# Conserved Pib2 regions have distinct roles in TORC1 regulation at the vacuole

**DOI:** 10.1101/2022.03.04.483060

**Authors:** Kayla K. Troutman, Natalia V. Varlakhanova, Bryan A. Tornabene, Rajesh Ramachandran, Marijn G.J. Ford

## Abstract

TORC1 is a critical controller of cell growth in eukaryotes. In yeast, the presence of nutrients is signaled to TORC1 by several upstream regulatory sensors that together coordinate TORC1 activity. TORC1 localizes to both vacuolar and endosomal membranes, where differential signaling occurs. This localization is mimicked by Pib2, a key upstream TORC1 regulator that is essential for TORC1 reactivation after nutrient starvation or pharmacological inhibition. Pib2 has both positive and negative effects on TORC1 activity, but the mechanisms remain poorly understood. Here, we pinpoint the Pib2 inhibitory function on TORC1 to residues within short, conserved N-terminal regions. We also show that Pib2 C-terminal regions, helical region E and tail, are essential for TORC1 reactivation. Further, the Pib2 FYVE domain is essential for vacuolar localization, however, it is surprisingly unnecessary for recovery from rapamycin exposure. Using chimeric Pib2 targeting constructs, we show that endosomal localization is not necessary for TORC1 reactivation and cell growth after rapamycin treatment. Thus, a comprehensive molecular dissection of Pib2 demonstrates that each of its conserved regions differentially contribute to Pib2 regulation of TORC1 activity.

## Introduction

The target of rapamycin complex I (TORC1) is a conserved kinase complex essential for the regulation of cell growth. In yeast, TORC1 responds to and integrates a diverse range of nutritional cues to promote anabolic processes such as protein synthesis, while inhibiting catabolic processes such as macroautophagy (hereafter autophagy). Inhibition of TORC1, pharmacologically or through nutrient starvation, triggers autophagy and reduces protein and ribosome synthesis (Gonzalez and Hall, 2017; Loewith and Hall, 2011). In metazoans, growth factor-mediated signaling pathways also converge on TORC1 to provide an additional layer of cell growth regulation. Given its central role in regulating cell growth, TORC1 has been implicated in a variety of diseases including cancer and diabetes, as well as aging (Gonzalez and Hall, 2017; Saxton and Sabatini, 2017).

Yeast TORC1 consists of two copies each of the following subunits: the serine/threonine kinase Tor1 (or Tor2), Kog1, Lst8, and Tco89 (Loewith et al., 2002; Reinke et al., 2004; Wedaman et al., 2003). TORC1 has several downstream effectors which regulate protein and ribosome synthesis and autophagy. Primary TORC1 effectors which promote protein synthesis include Sch9 and Rps6 (Gonzalez et al., 2015; Hatakeyama et al., 2019; Urban et al., 2007). Major effectors of autophagy include Atg13 and Vps27 (Hatakeyama et al., 2019; Kamada et al., 2010). TORC1 localizes to the vacuolar membrane as well as perivacuolar puncta (Binda et al., 2009; Kira et al., 2016; Sturgill et al., 2008). By colocalization with endosomal markers, these puncta have been identified as signaling endosomes where TORC1, along with its regulatory components and effectors colocalize (Chen et al., 2021; Hatakeyama et al., 2019). It has recently been reported that active vacuolar or endosomal TORC1 serve two different functions: where TORC1 activity at the vacuole results in phosphorylation of Sch9 to promote protein synthesis, and phosphorylation of Atg13 and Vps27 occurs at the endosomal pool of TORC1 to inhibit autophagy (Hatakeyama et al., 2019). Whether there are differences in the regulatory mechanisms at these distinct subcellular locations remains unclear.

In yeast, TORC1 is ultimately regulated by amino acids via several upstream proteins. A fundamental activator of TORC1 is the escape from rapamycin-induced growth arrest (EGO) complex (Peli-Gulli et al., 2015), which consists of the GTPases Gtr1 and Gtr2, and the docking proteins Ego1, Ego2, and Ego3 (Powis et al., 2015).Through the EGO complex, the amino acids glutamine and leucine are predominant activators of yeast TORC1 (Binda et al., 2009; Bonfils et al., 2012; Peli-Gulli et al., 2015). Several upstream GTPase-activating proteins (GAPs) and guanine nucleotide exchange factors (GEFs) control the nucleotide-binding status of the Gtr GTPases (Binda et al., 2009; Panchaud et al., 2013; Peli-Gulli et al., 2015). Together, under nutrient rich conditions, these GAPs and GEFs help to maintain an active EGO complex (wherein Gtr1 is GTP-bound and Gtr2 is GDP-bound) to promote TORC1 activation through direct interaction with TORC1 components (Binda et al., 2009; Gonzalez and Hall, 2017; Jeong et al., 2012; Nakashima et al., 1999; Panchaud et al., 2013; Peli-Gulli et al., 2015).

PI3P and Vps34, the kinase that synthesizes it, are also key TORC1 regulators. In mammalian cell models, this connection is well understood; amino acids activate the PI3K Vps34, which generates PI3P and results in mTORC1 activation through the Rag GTPases, the metazoan homologs of the Gtrs (Byfield et al., 2005; Nobukuni et al., 2005; Yoon et al., 2011). In yeast, Vps34 deletion strains show a loss of both PI3P and PI(3,5)P2 and are hypersensitive to the pharmacological TORC1 inhibitor, rapamycin (Bridges et al., 2012). PI3P is a precursor for the synthesis of PI(3,5)P2, which has also been shown to affect TORC1 localization and activity in both yeast and mammalian cells (Bridges et al., 2012; Chen et al., 2021). This highlights the role of these phosphoinositides in TORC1 regulation.

Phosphatidylinositol 3-phosphate-binding protein 2 (Pib2) is a master regulator of TORC1 signaling in yeast (Hatakeyama, 2021). Pib2 is unique in that it was identified in screens for both rapamycin sensitivity (Dubouloz et al., 2005; Parsons et al., 2004) and rapamycin resistance (Michel et al., 2017). Interestingly, Pib2 has two opposing functions in its regulation of TORC1: an inhibitory effect mediated by its N-terminal regions, and an activation effect mediated by its C-terminal domains (Michel et al., 2017; Sullivan et al., 2019). It has been demonstrated that Pib2 interacts with Tor1 and Kog1, as well as EGO complex components (Michel et al., 2017; Sullivan et al., 2019; Tanigawa and Maeda, 2017; Tarassov et al., 2008; Ukai et al., 2018). Pib2 is essential for reactivation of TORC1 following rapamycin exposure and in response to glutamine and leucine following nitrogen starvation (Varlakhanova et al., 2017). Recently, Pib2 has been implicated as a glutamine sensor which directly interacts with TORC1, in a glutamine-dependent manner, to promote TORC1 activity (Tanigawa and Maeda, 2017; Tanigawa et al., 2021; Ukai et al., 2018). However, the molecular mechanisms of Pib2’s regulatory interactions remain poorly understood.

Pib2 was named based on the presence of a C-terminal FYVE domain (Shin et al., 2001), a common structural module known to interact with PI3P, and it localizes to the vacuole and endosomes in a Vps34-dependent manner (Kim and Cunningham, 2015). In addition to the FYVE domain, Pib2 also contains a conserved and predicted alpha-helical region termed helical region E and a conserved C-terminal tail motif, as well as some shorter regions of conservation termed regions A-D, embedded in the otherwise low-complexity Pib2 N-terminal sequence. Pib2 induces lysosomal membrane permeabilization (LMP) and subsequent cell death through increased TORC1 activation in cells treated with ER stressors or calcineurin inhibitors (Kim and Cunningham, 2015). In this function, the C-terminal FYVE domain is essential for localization of Pib2 at vacuolar and endosomal membranes and both the FYVE domain and tail motif are necessary for TORC1 activation and promotion of LMP (Kim and Cunningham, 2015).

Here we performed a molecular dissection of Pib2 and show that each of its conserved regions differentially contribute to Pib2 localization and regulation of TORC1 activity. We demonstrate that conserved regions A and B within the Pib2 N-terminus are responsible for the inhibitory effect on TORC1 reactivation following rapamycin exposure, whereas the C-terminal alpha-helical E region and tail motif are essential for TORC1 reactivation. We assessed the role of Pib2 in TORC1 reactivation and cell growth at the endosome and the vacuole, showing that vacuolar localization of Pib2 is essential for recovery of cell growth. We also demonstrate that the alpha-helical E region and the FYVE domain play key roles in the vacuolar localization of Pib2. Further, the helical E region contains residues which are essential for TORC1 reactivation but not Pib2 localization, showing that vacuolar localization is necessary but not sufficient for TORC1 reactivation.

## Results

### Conserved Pib2 regions differentially regulate TORC1 reactivation

Previous studies have shown that the N- and C-terminal regions of Pib2 function in inhibiting and activating TORC1, respectively (Michel et al., 2017; Sullivan et al., 2019). We sought to identify the precise regions responsible for these dual functions. Through multiple sequence alignments (MSAs) using sequences of Pib2 homologs from 15 ascomycete fungi, seven conserved regions are apparent (Fig. 1A; Fig. S1). The regions identified include the N-terminal regions A-D, a FYVE domain, and a C-terminal tail motif. Previous alignments defined a smaller region E (Kim and Cunningham, 2015), which we extended to include other well-conserved residues (298-418). The AlphaFold2 prediction of Pib2 structural elements (Jumper et al., 2021), as well as other bioinformatic approaches, suggest this region has alpha-helical secondary structure, hence the name helical E region (helE) (Fig. 1A; Fig. S1C).

**Figure 1.**
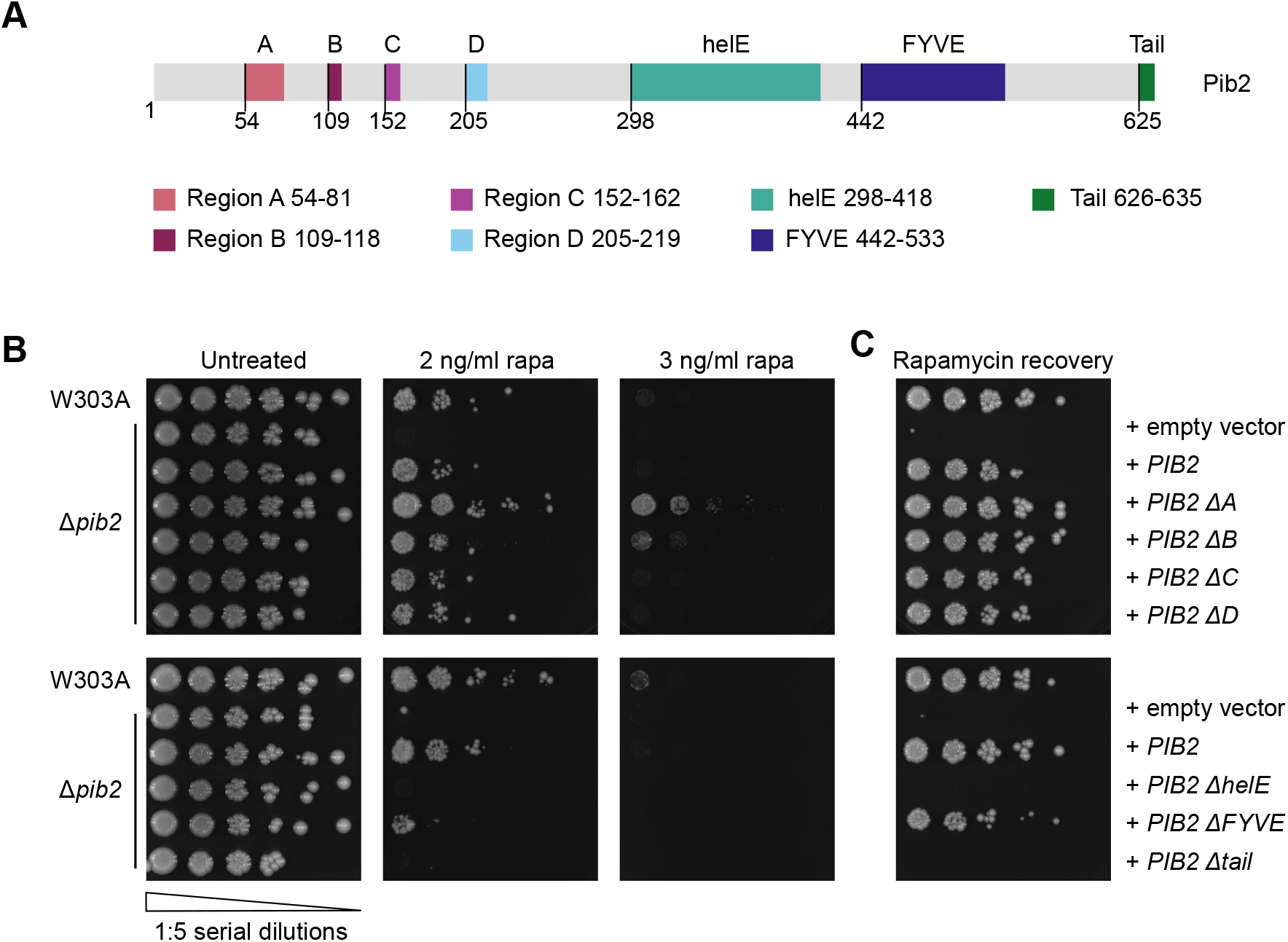
TORC1 reactivation is differentially regulated by conserved Pib2 regions. (A) The primary structure of Pib2, highlighting its conserved regions. The starting residue number of each region is shown below the schematic. Deletion constructs were generated by replacing the indicated regions with a short AGAGA linker. (B) Rapamycin exposure assay assessing growth of W303A and Δ*pib2* cells expressing the indicated constructs on YPD or YPD supplemented with the indicated concentration of rapamycin (rapa). The left-most spots correspond to 2 μl of a OD_600_ = 0.5 culture, followed by 2 μl of sequential 1:5 dilutions. (C) Rapamycin recovery assay assessing growth of W303A and Δ*pib2* expressing the indicated constructs on YPD following rapamycin exposure. Exponentially growing cells were treated with 200 ng/ml rapamycin in YPD at 30ºC for 4 hours. After washing, cells were plated on YPD and incubated at 30ºC for 3 days. Cells were plated as in B.

To elucidate the functions of these Pib2 regions, we generated deletion constructs for each of the 7 conserved Pib2 regions (Fig. 1A). Using Δ*pib2* cells and plasmid-expressed Pib2 region deletion constructs, we performed two types of rapamycin growth assays. We assessed growth of cells in the presence of a low concentration of rapamycin and on nutrient-rich plates following a 4-hour exposure to a high (200 ng/ml) concentration of rapamycin. These two approaches assay growth in an ongoing stress state and recovery from the stressor, respectively. Δ*pib2* cells expressing Pib2 ΔhelE and Pib2 ΔTail constructs did not grow on rapamycin-containing plates (Fig. 1B) and were unable to recover from rapamycin treatment (Fig. 1C). Unexpectedly, the Pib2 construct lacking its FYVE domain (Pib2 ΔFYVE) was able to grow on rapamycin-containing plates and recover from exposure to rapamycin, although at a much slower rate than isogenic wild-type (W303A) cells or Δ*pib2* cells expressing wild-type Pib2 (Fig. 1B-C). In contrast, deletion of Pib2 regions A or B resulted in enhanced growth in both rapamycin exposure assays (Fig. 1B-C). Deletion of Pib2 regions C or D did not influence cell growth on rapamycin plates or the ability to recover following rapamycin exposure (Fig. 1B-C). These results suggested that the N-terminal regions A and B were key for Pib2’s inhibitory function, whereas the C-terminal helE region and tail motif were essential for TORC1 reactivation. While previous studies used more extensive region deletions, our results are in agreement showing that the Pib2 N-terminal regions serve an inhibitory function, and the C-terminal regions are needed for TORC1 reactivation (Michel et al., 2017; Sullivan et al., 2019). As a control, the growth of the Pib2 deletion constructs did not differ on YPD plates without rapamycin exposure (Fig. 1B).

To ensure that the differences in rapamycin recovery were not due to changes in protein expression levels, we used Pib2 region deletion constructs tagged at the N-terminus with yEGFP and western blotting to confirm that these mutants are expressed at equal levels under logarithmic phase growth conditions (Fig. S2A-B). As a further control, we used GFP-Atg8 to assess the induction and flux of autophagy in response to both rapamycin exposure and nutrient starvation (Guan et al., 2001; Shintani and Klionsky, 2004). No differences were observed between W303A and Δ*pib2* cells in either condition (Varlakhanova et al., 2017) (Fig. S2C-D). Additionally, no increase in autophagy was observed in steady state in Δ*pib2* cells, thus Pib2 does not influence TORC1 repressive function on autophagy (Varlakhanova et al., 2017) (Fig. S2C-D). In addition, as the rapamycin assays used plasmid-based expression of Pib2, we generated genomic strains of *PIB2* ΔA, *PIB2* ΔB, and *PIB2* ΔhelE, as well as wild-type *PIB2*, where the only source of *PIB2* is its native genomic context, to demonstrate that plasmid expression was representative of endogenous expression. Using these genomically-modified strains, we repeated the rapamycin growth assays and observed that the genomic *PIB2* region deletion mutants showed the same growth patterns as the plasmid-expressed deletion constructs (Fig. S2E).

### Conserved Pib2 regions differentially affect Pib2 subcellular localization

TORC1 localizes to the both the vacuolar and endosomal membranes where differential TORC1 signaling occurs (Binda et al., 2009; Hatakeyama et al., 2019; Kira et al., 2016; Sturgill et al., 2008). As localization is likely an important aspect of Pib2’s ability to reactivate TORC1, we sought to determine how the Pib2 regions contribute to its subcellular localization. To this end, we generated the same Pib2 region deletion constructs N-terminally tagged with yEGFP. We first confirmed, using genomically yEGFP-tagged *PIB2*, that this tag does not interfere with Pib2’s ability to activate TORC1 and promote cell growth under these conditions (Fig. S3A). Thus, we used yEGFP-Pib2 region deletion constructs to assess localization of the Pib2 deletions via confocal microscopy (Fig. 2A-B). Wild-type Pib2 localized primarily to the vacuolar membrane with occasional perivacuolar puncta (Fig. 2A-C). This vacuolar localization phenotype was consistent for most of the Pib2 deletion constructs, however, we observed that deletion of the helE region or FYVE domain resulted in a mixed phenotype, with some vacuolar localization and a large cytosolic component (Fig. 2A-B). As both regions are necessary for vacuolar localization of Pib2, it suggests these domains may act as a potentially redundant dual recruitment mechanism.

**Figure 2.**
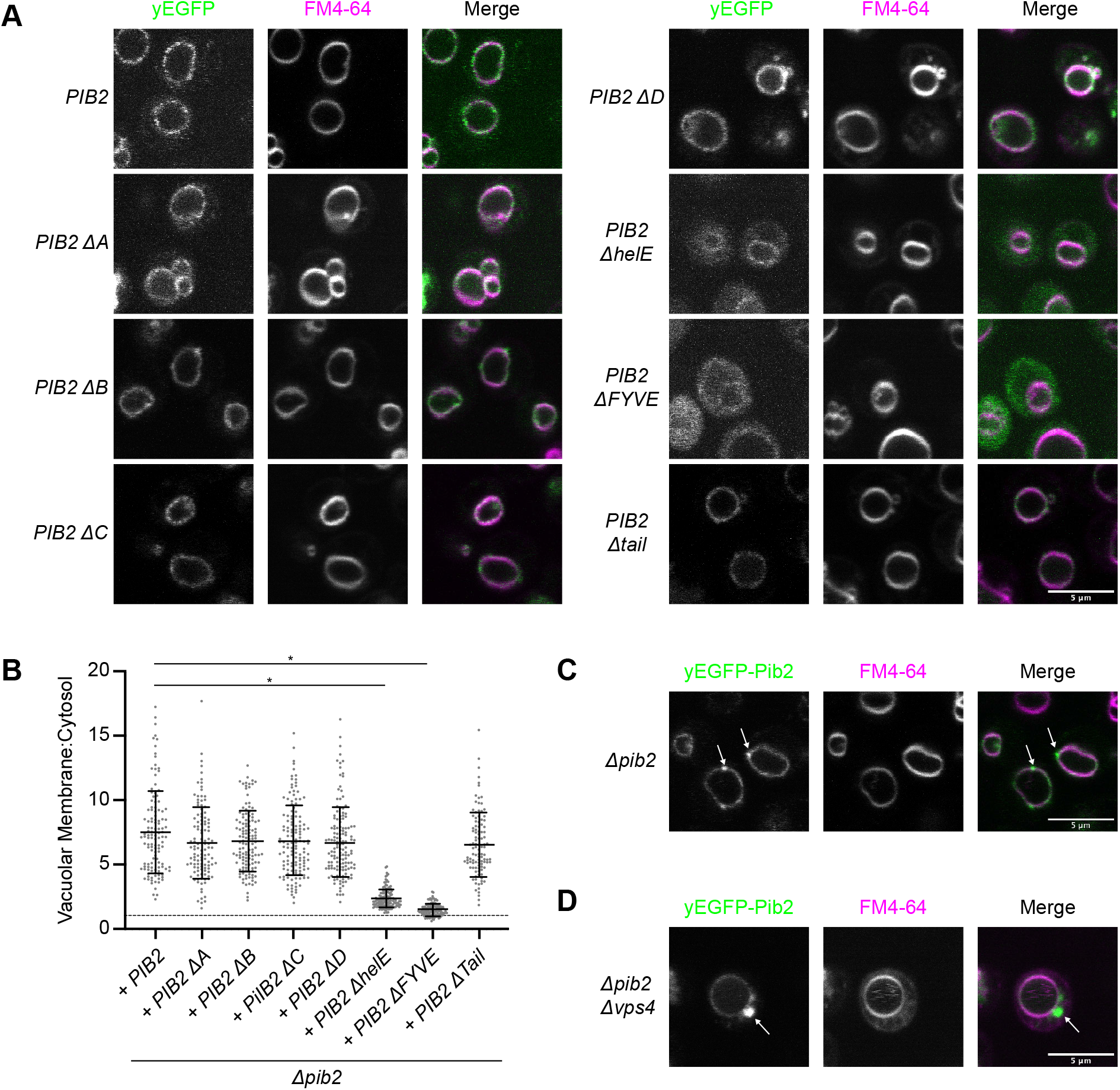
Pib2 localization is determined by its conserved regions. (A) Localization of Δ*pib2* cells expressing the indicated N-terminally yEGFP-tagged Pib2 constructs. For vacuolar visualization, cells were stained with FM4-64 prior to imaging. Representative fields of view are shown. (B) Quantification (mean±s.d.) of the data presented in A. Vacuolar localization is determined as the ratio of vacuolar yEGFP signal to cytosolic yEGFP signal. Following a ROUT outlier analysis (Q = 0.1%), a one-way ANOVA was conducted to determine differences in vacuolar localization (F = 121.6, P < 0.0001). Constructs significantly different from the localization of Pib2, as assessed by Tukey multiple post-hoc comparisons test, are indicated (*P < 0.0001). A total of 117-143 cells were quantified for each construct. (C) yEGFP-Pib2 expressed in a Δ*pib2* strain with perivacuolar puncta (arrows). (D) yEGFP-Pib2 expressed in Δ*pib2 Δvps4* cells showing Pib2 localization to the Class E compartment (arrow).

To further assess the subcellular localization of Pib2, we expressed yEGFP-Pib2 in a Δ*pib2* Δ*vps4* strain. Vps4 is an ATPase that is essential for endosomal morphology and endosome to vacuole transport (Babst et al., 1997; Babst et al., 1998). The vacuolar protein sorting (Vps) genes are divided into phenotypic classes based on the defects on vacuolar morphology caused by mutation and deletion of these genes (Coonrod and Stevens, 2010; Raymond et al., 1992). Vps4 is a Class E mutant; these mutants have enlarged prevacuolar endosomal structures called Class E compartments (Babst et al., 1997; Babst et al., 1998; Raymond et al., 1992). Using this deletion strain, we were able to show that the perivacuolar Pib2 puncta correspond to endosomal structures as evidenced by the presence of Pib2 on the Class E compartments (Fig. 2D). Vacuolar localization was not affected.

### N-terminal Pib2 regions A and B display a TORC1 inhibitory function

Previous studies have demonstrated a TORC1 inhibitory function for Pib2 as it pertains to TORC1 reactivation following rapamycin exposure (Michel et al., 2017). Large truncations of its N-terminal region, residues 1-164, have shown that in the absence of the N-terminus, Pib2 is better able to reactivate TORC1 and promote cell growth (Michel et al., 2017). In the rapamycin assays in Figure 1B, we demonstrated that Regions A (residues 54-81) and B (residues 109-118) were central to the Pib2 inhibitory function. As the Pib2 ΔA construct showed enhanced growth in the rapamycin exposure assays, we sought to determine if the increased growth rate and TORC1 activity is specific to TORC1 reactivation. So, we assessed growth of select Pib2 constructs in liquid culture. In nutrient replete growth conditions (YPD), we found that the Pib2 ΔA construct grew at the same rate at wild-type Pib2 (Fig. S3B). This suggests the difference in growth rates between WT Pib2 and Pib2 ΔA, as observed in rapamycin exposure assays is specific to TORC1 reactivation. Further, we expressed the Pib2 ΔA construct in a Δ*gtr1*/*gtr2* strain. Pib2 ΔA was unable to rescue the rapamycin sensitivity of this strain, as it did not allow for growth on rapamycin plates or recovery from rapamycin exposure (Fig. S3C). To further uncover the inhibitory mechanism of Pib2 we used our MSAs to select highly conserved residues within regions A and B which might be involved in TORC1 inhibition. In region A we identified a series of conserved lysines, lysines 59-61 (Fig. 3A). As these lysines would be susceptible to post-translational modifications we generated a lysine to alanine mutant, Pib2 KA. Pib2 KA mutant grew on rapamycin-containing plates and recovered from rapamycin exposure at a similar rate to the Pib2 ΔA construct and did not affect Pib2 subcellular localization (Fig. 3B-D). Further, as the positive charge of these lysines could form part of a binding interface and result in interactions with negatively charged residues in a binding partner, we also mutated these residues to glutamates (Pib2 KE). This mutant also grew at a similar rate to the Pib2 ΔA and Pib2 KA mutants (Fig. 3B). As mutation from lysine to either alanine or glutamate had a similar effect, we propose that the positive charge of the lysines in this region are key to its TORC1 inhibitory function.

**Figure 3.**
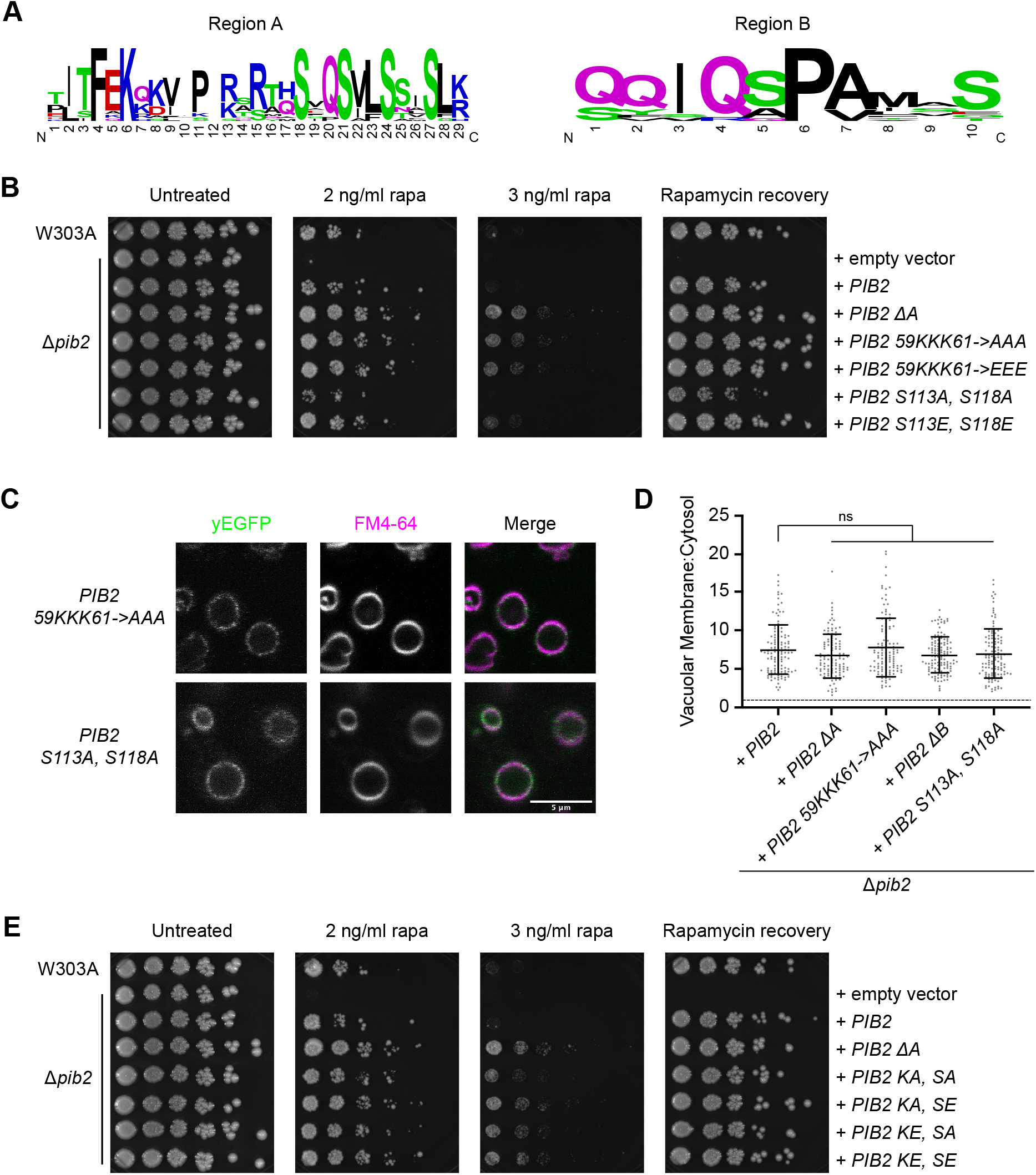
Key residues in Pib2 regions A and B are involved in the TORC1 inhibitory mechanism. (A) Sequence logos illustrating the residues conserved in Pib2 regions A and B (15 ascomycete fungi sequences used for alignment). Residue label size is proportional to conservation. (B) Rapamycin exposure and recovery assays of Δ*pib2* cells expressing the indicated constructs. These were performed as in Figure 1. (C) Vacuolar localization of indicated yEGFP-Pib2 mutant constructs. Vacuoles were stained with FM4-64. (D) Quantification (mean±s.d.) of the data presented in C. Data for Pib2, Pib2 ΔA, and Pib2 ΔB were the same as in Figure 2B. Following a ROUT outlier analysis (Q = 0.1%), a one-way ANOVA was conducted to determine differences in vacuolar localization (F = 3.040, P = 0.0169). There were no significant differences from Pib2 localization as determined by Tukey multiple comparisons test. A total of 117-126 cells was quantified for each construct. (E) Rapamycin exposure and recovery assays of Δ*pib2* cells expressing the indicated constructs. These were performed as in Figure 1.

In region B, we identified conserved serines S113 and S118. Bioinformatic analyses of the Pib2 sequence showed that S113 is a predicted phosphorylation site (Ingrell et al., 2007). We therefore assessed phospho-dead and phosphomimetic mutants for growth on rapamycin plates and recovery from rapamycin exposure. The phospho-dead mutant, Pib2 S113A, S118A (Pib2 SA), showed reduced growth compared to wild-type Pib2 and did maintain vacuolar localization (Fig. 3B-D). However, the phosphomimetic mutant, Pib2 S113E, S118E (Pib2 SE), grew better than wild-type Pib2 in these rapamycin exposure assays (Fig. 3B). This is most notable on the 2 ng/ml rapamycin plate and the effects are diminished with increased rapamycin concentration (Fig. 3B). This suggests that phosphorylation at one or both of these sites may be involved in Pib2’s inhibitory mechanism.

A potential model of TORC1 inhibition by Pib2 could involve intramolecular interactions. To determine if regions A and B work together to inhibit TORC1 activity, we generated 4 combinatorial mutants of these key region A and B residues. These included: Pib2 KA/SA, Pib2 KA/SE, Pib2 KE/SA, and Pib2 KE/SE. Each of these mutants showed improved growth over wild type in both rapamycin exposure assays (Fig. 3E). As the lysine mutations in region A were able to override the activity of the region B serine mutations, this suggests region A is dominant to region B in this mechanism. To determine if Pib2 inhibitory regions interact with Pib2 C-terminal activation regions we generated compound deletion mutations. Deletion of region A in combination with one of the C-terminal activation regions (helical E, FYVE, or tail) enabled mild recovery following rapamycin exposure compared to deletion of the C-terminal regions alone (Fig. S3D). While this was a less robust recovery, this indicates that Pib2’s N- and C-terminal mechanisms of inhibition and activation of TORC1 may be separable.

### Pib2 vacuolar localization is essential for TORC1 reactivation and cell growth

Pib2 localizes to both the vacuole and endosomes (Hatakeyama et al., 2019; Kim and Cunningham, 2015). As Pib2 localization is similar to that of TORC1, we set out to investigate which cellular localization of Pib2 is essential for TORC1 reactivation. We therefore generated several chimeric Pib2 constructs targeted to the vacuole and endosome. These constructs consisted of a targeting protein linked to Pib2 via a yEGFP bridge (Fig. 4A). To target Pib2 specifically to the vacuole we used Vac8, which is localized to the vacuole by N-terminal lipid modifications (myristoylation and palmitoylation) (Subramanian et al., 2006). To target Pib2 specifically to endosomes we used the yeast SNX-BAR, Mvp1, which uses its PX and BAR domains to specifically bind endosomal membranes, which are enriched in PI3P (Sun et al., 2020) (Fig. 4A). The Vac8-yEGFP-Pib2 construct localized to the vacuolar membrane with scarce endosomal puncta (Fig. 4B, S4B). In contrast, the Mvp1-yEGFP-Pib2 construct localized primarily to endosomal puncta with minimal vacuolar localization (Fig. 4B, S4B). In response to rapamycin exposure assays, cells containing Vac8-yEGFP-Pib2 were able to grow on rapamycin containing plates and recover from rapamycin exposure, as expected (Fig. 4C). Growth of cells containing Mvp1-yEGFP-Pib2, however, was significantly compromised in response to rapamycin exposure (Fig. 4C).

**Figure 4.**
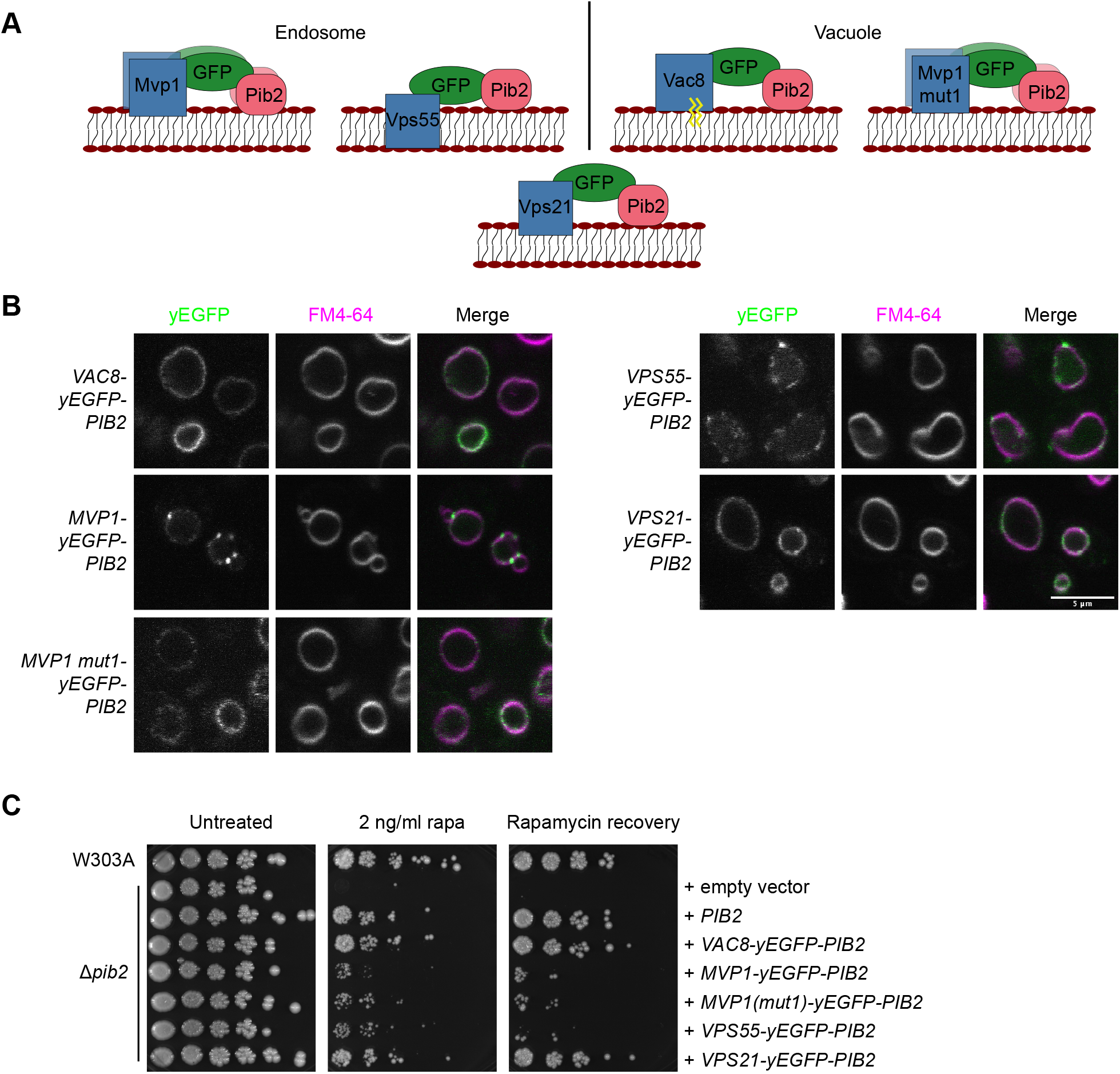
Pib2 vacuolar targeting promotes TORC1 reactivation. (A) Schematics of chimeric Pib2 vacuolar and endosomal targeting constructs. Each construct consists of the targeting protein and yEGFP fused to the N-terminus of Pib2. (B) Localization of the indicated targeting constructs. Vacuoles were stained with FM4-64. (C) Rapamycin exposure and recovery assay of Δ*pib2* cells expressing the indicated targeting constructs. These were performed as in Figure 1.

Pib2 has been suggested to be a glutamine sensor (Tanigawa and Maeda, 2017; Tanigawa et al., 2021; Ukai et al., 2018). Thus, we additionally assessed cell growth after nutrient starvation and stimulation with glutamine using phosphorylation of Rps6 at S232/S233 as a readout of TORC1 activity (Gonzalez et al., 2015). We demonstrated that the endosomal Mvp1-yEGFP-Pib2 construct is deficient in reactivating TORC1 signaling following nitrogen starvation and glutamine stimulation compared to the vacuolar Vac8-yEGFP-Pib2 construct and WT Pib2 (Fig. S4C-D). To confirm that all the targeting constructs were expressed at similar levels, we quantified total GFP fluorescence in cells expressing these constructs and compared it to the fluorescence from cells expressing only yEGFP-Pib2 (Fig. S4A). This method was used due to concerns over western blotting transfer efficiency with these large constructs of differing size.

Further, we generated an Mvp1 mutant targeting construct, which will be referred to as Mvp1 mut1. We have previously shown that this mutation results in cytosolic localization of Mvp1 (Sun et al., 2020). This mutation allowed for Pib2’s regions and domains to direct the localization of the construct and resulted in a subcellular distribution much closer to that of wild-type Pib2 (Fig. 4B). However, this mutant was still unable to rescue growth under rapamycin exposure conditions (Fig. 4C). As both Mvp1 and Mvp1 mut1 are known to multimerize (tetramer and dimer, respectively - (Sun et al., 2020)), we reason that this may preclude Pib2 from reactivation of TORC1 in these conditions. Thus, we generated two additional targeting constructs to assess TORC1 reactivation. These included Vps55-yEGFP-Pib2 and Vps21-yEGFP-Pib2. Vps21-yEGFP-Pib2 localized to the vacuole with few perivacuolar puncta (Fig. 4B, S4B). Conversely, Vps55, an integral membrane protein involved in endosome to vacuole trafficking (Belgareh-Touze et al., 2002), localized primarily to perivacuolar puncta with less vacuolar presence (Fig. 4B, S4B). Like the Mvp1-yEGFP-Pib2 construct, Vps55-yEGFP-Pib2 was unable to rescue growth in the rapamycin exposure assays (Fig. 4C). The vacuolar Vps21-yEGFP-Pib2 however, was able to rescue growth in these assays (Fig. 4C). This suggests that vacuolar Pib2 is necessary for the reactivation of TORC1 and resumed cell growth following rapamycin exposure.

### Pib2 vacuolar localization is dependent on its helE and FYVE domains

The vacuolar localization of Pib2 has been shown to depend on PI3P and Vps34 (Kim and Cunningham, 2015). PI3P binding motifs within FYVE domains include WxxD and R+HHCRxCG (where x is any amino acid) (Burd and Emr, 1998; Stenmark et al., 2002). As these are both conserved across Pib2 FYVE domains (Fig. 5A, S1A), we generated two mutants, one for each of these motifs: Pib2 W449A, D452A (Pib2 WD) and Pib2 R470A, H472A, H473A (Pib2 RHH) (Fig. 5A). Both mutants showed compromised localization and were primarily cytosolic (Fig. 5B-C). Mutation of 3 highly conserved residues in helE, 339VLR341→AAA (Pib2 VLR), did not influence vacuolar localization (Fig. 5A-C). It remains unclear what aspect of the helE region is needed for vacuolar localization. Simultaneous deletion of both the helE region and FYVE domain resulted in a primarily cytosolic phenotype as expected (Fig. 5B-C).

**Figure 5.**
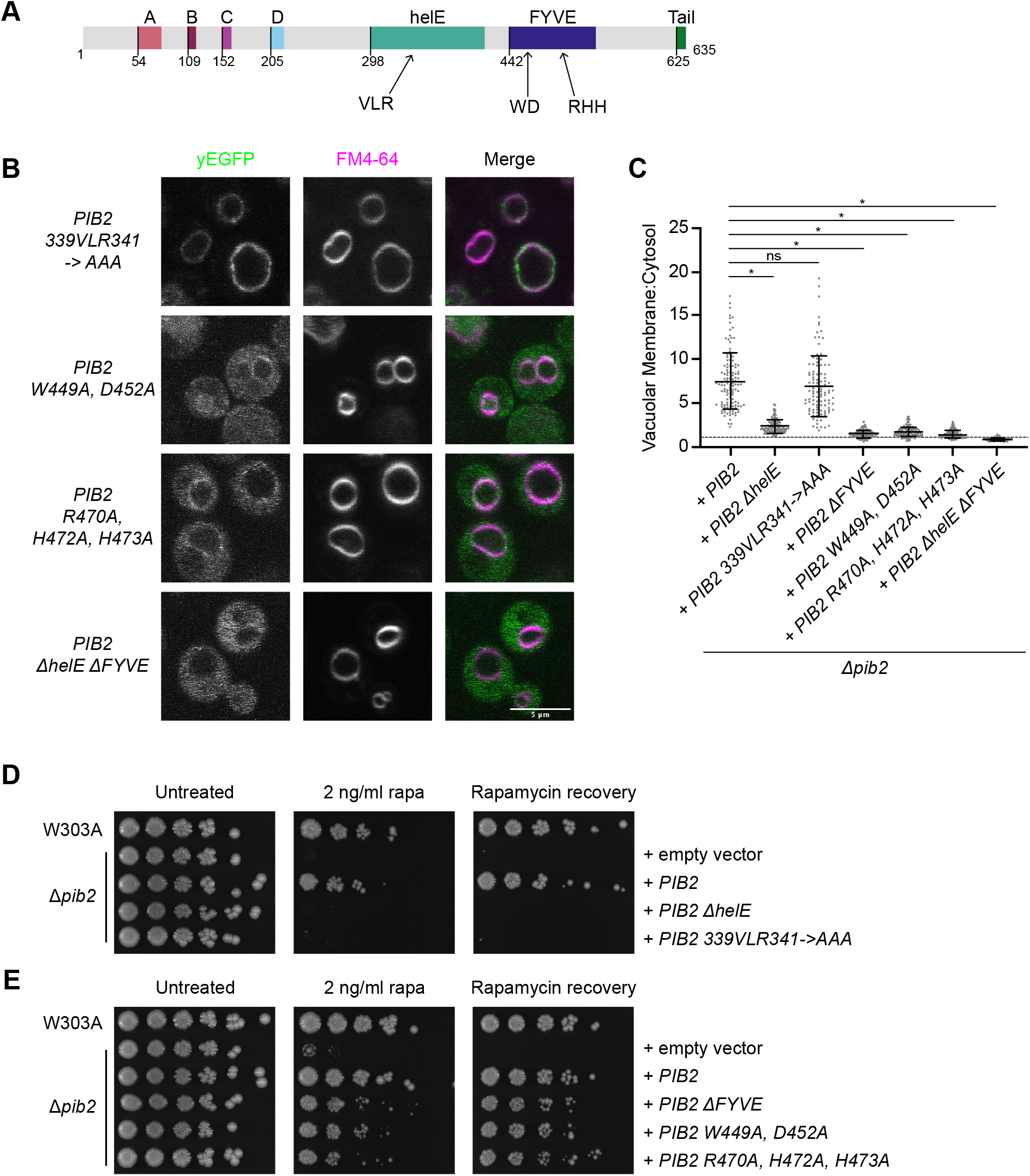
Pib2 helE region and FYVE domain are essential for TORC1 reactivation. (A) Schematic of Pib2 conserved regions with key helE region and FYVE domain residues indicated. (B) Localization of the indicated yEGFP-Pib2 mutants. Vacuoles were stained with FM4-64. (C) Quantification (mean±s.d.) of the data presented in B. Data for Pib2, Pib2 ΔhelE, and Pib2 ΔFYVE were the same as in Figure 2B. Following a ROUT outlier analysis (Q = 0.1%), a one-way ANOVA was conducted to determine differences in vacuolar localization (F = 294.8, P < 0.0001). Constructs significantly different from WT Pib2, as assessed by Tukey post-hoc multiple comparisons test, are indicated (*P < 0.0001). A total of 107-131 cells were quantified for each construct. (D) Rapamycin exposure and recovery assays of Δ*pib2* cells expressing the indicated Pib2 helE region mutants. These were performed as in Figure 1. (E) Rapamycin exposure and recovery assays of Δ*pib2* cells expressing the indicated Pib2 FYVE domain mutants. These were performed as in Figure 1.

As the FYVE domain alone was not sufficient to direct Pib2 localization to the vacuole, we investigated the ability of the Pib2 FYVE domain to bind to PI3P using isothermal titration calorimetry (ITC). We expressed and purified Pib2 442-625, which included the start of the FYVE domain to the beginning of the tail motif (Fig. S5A). ITC with Pib2 442-625 showed a weak binding interaction with the PI3P headgroup, inositol 1,3 bisphosphate, with a mean K_D_ = 212 μM, as well as the short chain lipid, PI3P diC4 (mean K_D_ = 736 μM) (Fig. S5B, Table S1). A mutant version of this construct, Pib2 442-625 RHH, did not bind to the lipid headgroup, as expected (Table S1). While many FYVE domains are monomeric, some are known to dimerize to facilitate PI3P binding (Kutateladze et al., 1999), so we additionally used size exclusion chromatography coupled with multi-angle light scattering (SEC-MALS) to determine the oligomeric state of the purified FYVE domain. Pib2 442-625 was predominantly monomeric (Fig. S5C). As this construct was much larger than the FYVE domain alone, we expressed and purified a smaller construct, Pib2 437-542. SEC-MALS with this construct again showed a largely monomeric population (Fig. S5D). As the binding affinity was relatively weak, it further supports that both the helE and FYVE domain are needed for appropriate subcellular localization of Pib2.

### Pib2 C-terminal regions are essential for TORC1 reactivation following rapamycin exposure

Previous studies have shown that truncation of the Pib2 C-terminal tail, and larger C-terminal portions, impedes the reactivation of TORC1 following rapamycin exposure (Michel et al., 2017; Ukai et al., 2018). As shown in Figure 1B-C, helE and tail motif were essential for TORC1 reactivation under rapamycin exposure conditions. The FYVE domain also influenced TORC1 reactivation, although less so than the other two domains. To determine what aspects of these conserved regions may be key for TORC1 reactivation, we assessed the sequence alignments for highly conserved residues. We then generated point mutations within helE and FYVE domain to assess their effects on TORC1 activation. The helE region mutant Pib2 VLR was not able to grow on rapamycin plates and did not recover from rapamycin exposure, similar to the helE region deletion construct (Fig. 5D). However, unlike the helE region deletion construct, Pib2 VLR localized predominately to the vacuole, resembling WT Pib2 localization (Fig. 5B-C). The helE region of Pib2 is a putative Kog1 binding region (Michel et al., 2017; Sullivan et al., 2019), thus it is possible that these residues may be required for that interaction.

We also assessed growth of the PI3P binding motif mutations within the Pib2 FYVE domain, Pib2 WD and Pib2 RHH. In addition to their altered localization described above (Fig. 5B-C), these mutants phenocopied deletion of the FYVE domain in that they recovered at a slower rate than wild-type Pib2 (Fig. 5E).

To determine if vacuolar localization of Pib2 was sufficient for TORC1 reactivation, we incorporated two of the previously described Pib2 mutations, Pib2 VLR and Pib2 WD, into the Vac8-yEGFP-Pib2 construct. As expected, due to the presence of Vac8, these mutants localized to the vacuole (Fig. 6A). The Vac8-yEGFP-Pib2 WD mutant was able to rescue growth under rapamycin exposure conditions just as well as the WT Vac8-yEGFP-Pib2 construct (Fig. 6B). As vacuolar localization of the Pib2 WD mutant with this targeting construct was able to rescue growth rate, this suggests that vacuolar localization of Pib2 is necessary for TORC1 reactivation in these conditions. The Vac8-yEGFP-Pib2 VLR mutant, however, was still unable to rescue growth (Fig. 6B). These results support the hypothesis that these VLR residues within the helE region of Pib2 are essential for TORC1 reactivation and may be involved in glutamine sensing (Tanigawa et al., 2021) and/or direct interactions with TORC1 components. Overall, these results demonstrate that vacuolar localization of Pib2 is necessary but not sufficient for TORC1 reactivation following rapamycin exposure in vivo.

**Figure 6.**
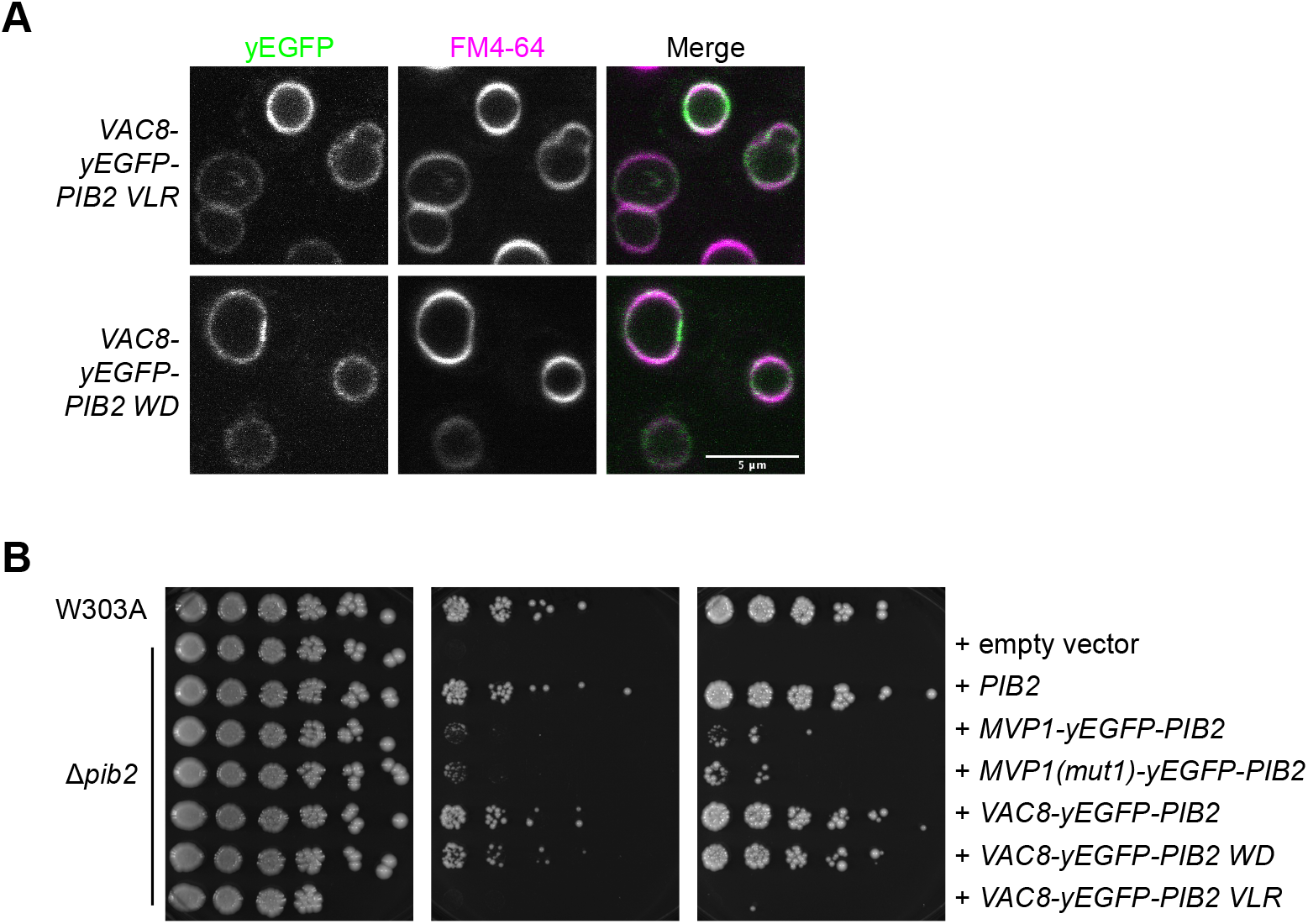
Vacuolar localization of Pib2 is necessary but not sufficient for TORC1 reactivation. (A) Localization of the indicated vacuolar targeting constructs. Vacuoles stained with FM4-64. (B) Rapamycin exposure and recovery assays of Δ*pib2* cells expressing the indicated targeting constructs. These were performed as in Figure 1.

**Figure 7.**
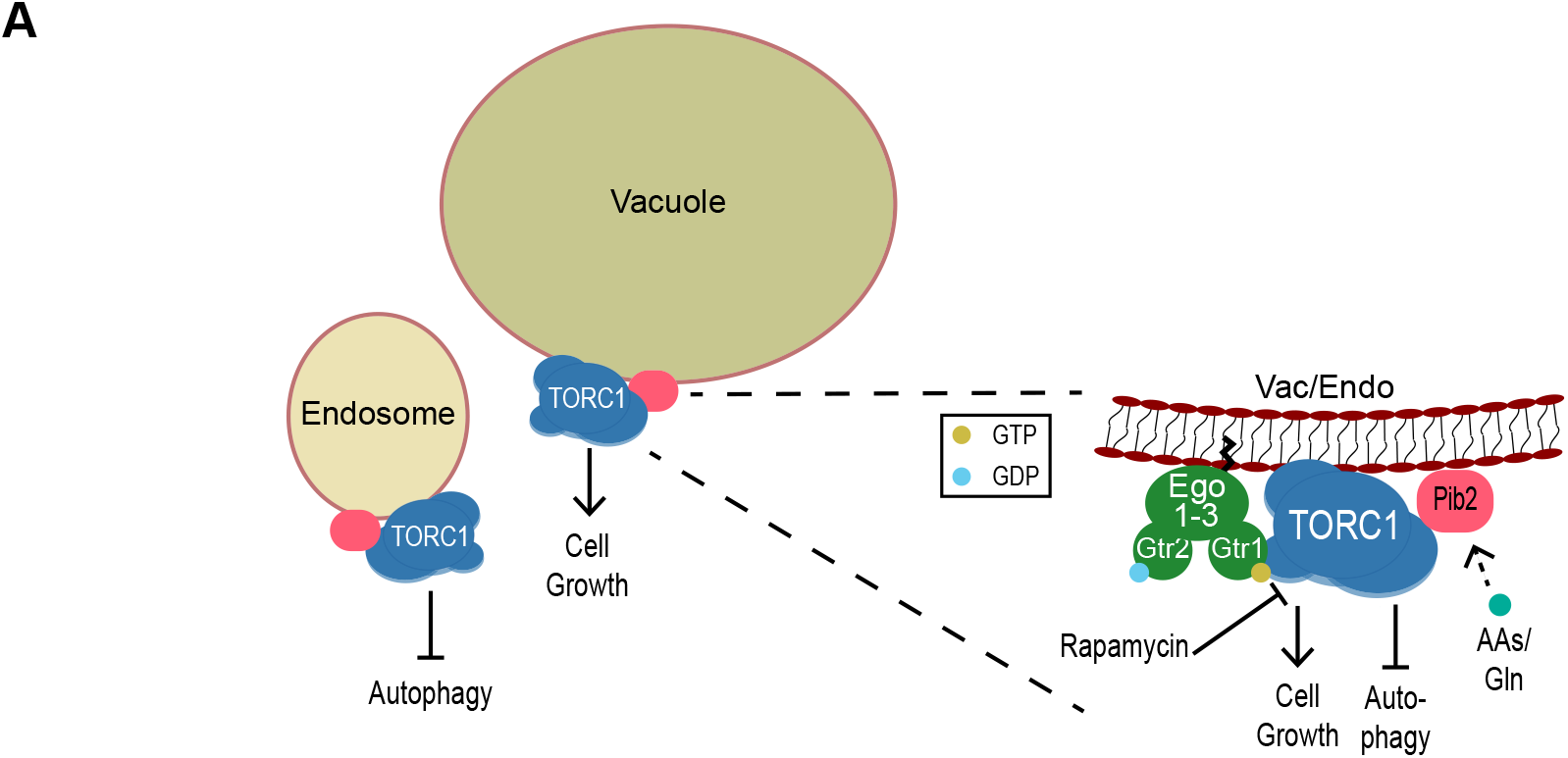
Proposed model of Pib2 regulation of TORC1.

## Discussion

In this work, we explored the roles of the conserved Pib2 regions in regulating TORC1 activity. We demonstrated that each conserved region has a distinct function in TORC1 reactivation or inhibition, and/or Pib2 subcellular localization. Pib2 N-terminal regions A and B displayed a TORC1 inhibitory function, as demonstrated by enhanced growth of Pib2 ΔA, Pib2 ΔB, and related mutants over wild-type Pib2 in rapamycin exposure assays. C-terminal Pib2 regions, notably helE and the tail motif were essential for TORC1 reactivation, as demonstrated by the lack of growth and recovery in rapamycin exposure assays. Further, the C-terminal helE and FYVE regions were key for vacuolar localization of Pib2, and this localization was imperative for its function. Interestingly, while the Pib2 ΔFYVE construct was largely cytosolic, it was still able to reactivate TORC1 in rapamycin assays, although at a slower rate than wild-type Pib2. We speculate that this could be due to residual Pib2 localization at the vacuole mediated by the helical E region which allows for some interaction with TORC1 to occur.

Pib2 C-terminal regions have been shown to be essential for TORC1 reactivation. Deletion mutants of the tail motif or Kog1 binding domains (found within the conserved helE region) exhibit increased rapamycin sensitivity (Michel et al., 2017; Sullivan et al., 2019). We demonstrated here that helE specific VLR sequence and tail motif were required for Pib2’s ability to grow on rapamycin-containing plates and recover following rapamycin exposure (Fig. 1B-C). The mechanism of the tail motif’s involvement in TORC1 reactivation is unclear. However, several hyperactive tail mutations were recently reported and it was proposed that the tail directly interacts with TORC1 to promote its activation (Tanigawa et al., 2021).

Pib2 interaction with TORC1 components, Kog1 and Tor1, have previously been demonstrated by western blotting and yeast two-hybrid experiments (Michel et al., 2017; Sullivan et al., 2019; Tanigawa and Maeda, 2017; Ukai et al., 2018). The helE region has been identified as a putative binding region for these interactions (Michel et al., 2017; Sullivan et al., 2019). As confocal imaging of our Pib2 ΔhelE mutant showed, this region is partially responsible for Pib2 localization at the vacuole (Fig. 2A-B). However, the helE mutant Pib2 339VLR341−>AAA demonstrated that these residues were essential for TORC1 reactivation following rapamycin exposure, even though they have no effect on vacuolar localization. It has recently been reported that these residues, particularly R341, may be essential for glutamine-induced TORC1 activation (Tanigawa et al., 2021), which could be a part of the TORC1 interaction and activation mechanism. However, further experiments are needed to define how Pib2 interacts with TORC1 components or other TORC1 regulators. One potential model is that these residues are crucial for the interaction between Pib2 and TORC1 components needed for TORC1 reactivation. We were unable to reliably detect an interaction between WT Pib2 and Kog1 using yeast two-hybrid experiments, so potential effects of the 339VLR341−>AAA mutation on this interaction could not be confirmed.

Previous studies have shown that truncations of the Pib2 N-terminus result in increased resistance to rapamycin exposure (Michel et al., 2017; Ukai et al., 2018), suggesting these regions have inhibitory effects on TORC1 reactivation. Our results showed that regions A and B are required for this inhibitory role. In contrast, deletion of regions C and D had no effect on TORC1 reactivation with the readouts used here. Pib2 MSAs showed fewer conserved residues in these regions compared to regions A and B so they may not be vital for this function (Fig. S1A). Regions C and D could therefore play a role in other Pib2 functions, such as LMP (Kim and Cunningham, 2015). Mutation of conserved residues within regions A and B highlighted a series of lysines and serines involved in the inhibitory mechanism. Based on our results and bioinformatic post-translational modification (PTM) predictions, the positive charge of the lysines and the phosphorylation status of the serines are likely a key part of this mechanism. Other groups have reported phosphorylation of Pib2 by the TORC1 downstream kinase Npr1 (Brito et al., 2019; MacGurn et al., 2011) and there are several predicted phosphorylation sites within the Pib2 N-terminus which could affect its function (Fig. S1A). Another potential kinase candidate includes Cdk1, which has previously been shown to phosphorylate Pib2 (Holt et al., 2009). As the AlphaFold2 prediction shows, these N-terminal regions are found in low complexity sequence, which not only makes them more accessible and susceptible to PTMs but also protein-interactions which could play a role in TORC1 regulation. Further studies are needed to elucidate this inhibitory mechanism.

The Gtr-dependent pathway of TORC1 regulation relies on nutrient sensing by upstream GAPs and GEFs to control TORC1 activation. Whether Pib2 activation of TORC1 is a Gtr-dependent or Gtr-independent mechanism remains uncertain. Simultaneous deletion of Pib2 and EGO components results in synthetic lethality (Kim and Cunningham, 2015). Further, Pib2 interactions with EGO components have been shown by yeast two-hybrid and both Pib2 and EGO have been shown to be required for TORC1 activation by amino acids (Kim and Cunningham, 2015; Tarassov et al., 2008; Varlakhanova et al., 2017). However, some studies have described a parallel, Gtr-independent activation pathway via glutamine (Tanigawa and Maeda, 2017; Tanigawa et al., 2021; Ukai et al., 2018). Expression of our Pib2 ΔA construct, which confers rapamycin resistance, in a Δ*gtr1*/*gtr2* yeast strain did not allow for growth on rapamycin plates or recovery from rapamycin exposure (Fig. S3C), supporting that Pib2 and the Gtrs may work in the same pathway to promote TORC1 reactivation. A model of dual-phase activation has been suggested in which Pib2 and the Gtrs work cooperatively in some instances and independently in others to promote TORC1 activity (Hatakeyama, 2021). As it is possible to generate both Δ*pib2* and Δ*gtr1*/*gtr2* strains independently, it implies that under nutrient replete, unstressed conditions both pathways are not necessary for TORC1 activity. However, the inability of these strains to recover from stressors like rapamycin suggests that under stress both the Pib2 and Gtr pathways may be required for TORC1 activation.

Subcellular localization has shown to be an important aspect of TORC1 regulation and activity (Hatakeyama et al., 2019; Prouteau et al., 2017). Recently, two spatially distinct pools of TORC1 have been identified that direct different processes, with vacuolar TORC1 promoting cell growth and protein synthesis, and endosomal TORC1 negatively regulating autophagy (Hatakeyama et al., 2019). The Pib2 FYVE domain has been shown to be key for Pib2 localization at the vacuolar membrane (Kim and Cunningham, 2015). FYVE domains are most notably known as PI3P-binding domains and typically display endosomal localization (Stenmark et al., 2002). As that is not the case with Pib2, it is possible that the Pib2 FYVE domain may preferentially bind other lipids to help direct it to the vacuole. Indeed, our measurements indicate weak affinities for both the headgroup and a soluble PI3P. As purified FYVE constructs are predominately monomeric, regulation of assembly into a dimeric form may also be required to facilitate interaction with its preferred ligand. Further, we show here that the FYVE domain alone is not sufficient for Pib2 vacuolar localization. Our data showed that while the Pib2 ΔFYVE construct had a mostly cytosolic cellular distribution, it did still have some enrichment at the vacuolar membrane and still enabled recovery from rapamycin exposure, albeit at a slower rate (Fig. 1B-C, 2A-B). The Pib2 helE deletion construct, like the FYVE deletion construct, showed a mostly cytosolic phenotype with some vacuolar enrichment, however, it did not allow for recovery from rapamycin exposure (Fig. 1B-C, 2A-B). Here, we used endosomal and vacuolar targeting constructs to identify the role of Pib2 at these subcellular localizations. The Vps55-yEGFP-Pib2 construct, that has an endosomal distribution, showed that endosomal Pib2 is not sufficient for TORC1 reactivation and cell growth. Further, vacuolar targeting by the Vac8-yEGFP-Pib2 WD construct rescued the growth deficit seen with the largely cytosolic Pib2 WD mutant. This suggests vacuolar localization of Pib2 is essential for cell growth and recovery following rapamycin exposure. While vacuolar localization of Pib2 appears to be essential for cell growth following rapamycin exposure, it is not sufficient to promote TORC1 activity. The helE region VLR mutant, which maintains vacuolar localization of Pib2, still did not support TORC1 reactivation (Fig. 5D, 6B). This implies that other interactions at the vacuolar membrane are necessary for Pib2 to reactivate TORC1. Future experiments could use a light-inducible dimerization system to alter the distribution of Pib2 or other TORC1 regulators and assess the role of Pib2 localization in TORC1 activation.

In summary, we have shown that Pib2 has two opposing functions in TORC1 regulation, and each conserved region plays a distinct role in inhibiting or reactivating TORC1. Additionally, using endosomal and vacuolar targeting constructs, we have shown that Pib2 localization at the vacuole is essential but not sufficient for promoting TORC1 activity and cell growth. We propose that an interaction between TORC1 and Pib2 at the vacuole is necessary for TORC1 reactivation and cell growth following exposure to a stressor. Future work will investigate the mechanisms of TORC1 inhibition by Pib2, as well as the implications of Pib2’s localization and dependence on other TORC1 regulatory proteins.

## Materials and Methods

### Multiple Sequence Alignments (MSA)

Sequence alignments were made using MUSCLE (Edgar, 2004). Alignments shown were made using the output of ESPRIPT 3.0 (Robert and Gouet, 2014) using the AlphaFold2 Pib2 prediction (PDB AF-P53191-F1-model_v1) (Jumper et al., 2021) as an input for secondary structure assignments. The AlphaFold2 structure prediction was rendered using PyMOL (Schrodinger, 2015).

### Media

YPD (1% yeast extract, 2% peptone, 2% glucose, supplemented with L-tryptophan and adenine) was used for routine growth. Synthetic complete (SC; yeast nitrogen base, ammonium sulfate, 2% glucose, amino acids) or synthetic defined (SD; yeast nitrogen base, ammonium sulfate, 2% glucose, appropriate amino acid dropout) media were used as indicated prior to microscopy or to maintain plasmid selection. For sporulation, cells were successively cultured in YPA (1% yeast extract, 2% peptone, 2% potassium acetate) and SPO (1% potassium acetate, 0.1% yeast extract, 0.05% glucose). For nitrogen starvation, cells were grown in SD–N (0.17% yeast nitrogen base without amino acids and ammonium sulfate, 2% glucose).

### Yeast genetic manipulation and cloning

Yeast strains used in this work are listed in Supplemental Table S2. W303A GFP-N10-*PIB2*::HIS3 (PY_262) was generated by complete replacement of the kanMX cassette in W303A Δ*pib2*::KAN (PY_126) (Varlakhanova et al., 2017). The replacement cassette, which includes ~40 nucleotide regions from the *PIB2* promoter and terminator sequences immediately upstream and downstream of the *PIB2* ATG and STOP codons to enable site-specific reintegration, was generated by splicing by overlap extension PCR using a GFP-N10-*PIB2* fragment amplified from pRS316 yEGFP-N10-*PIB2* + UTRs (Varlakhanova et al., 2017) and an ADH1 terminator-His3MX6 fragment, amplified from pFA6a-link-yEGFP-SpHIS5 (pKT0128) (Sheff and Thorn, 2004). The full-length replacement cassette was gel purified and transformed into PY_126. Cells were plated onto SD-HIS and incubated for 3 days at 30 °C. A mixture of recombinants was expected: replacement of the kan resistance gene by conversion to HIS prototrophy, without integration of GFP-N10-*PIB2* (due to recombination between the *Ashbya gossipii* TEF promoter and terminators sequences flanking the kanamycin resistance gene in Δ*pib2*::KAN and the *S pombe* his5 gene in the replacement cassette) and complete replacement of kanMX6 with the GFP-N10-*PIB2*::HIS replacement cassette, by recombination in the *PIB2* promoter and terminator regions. Colony PCR indicated ~1/3 of the resulting HIS prototrophs contained the GFP-N10-*PIB2* coding sequence at the correct genomic locus. Loss of kanamycin resistance was also verified. Candidate colonies were backcrossed to W303α and re-sporulated to ensure the expected pattern of segregation. The knock-in strain was further verified by sequencing, imaging and functional assays.

W303A *PIB2*::KAN, W303A *PIB2* ΔA::KAN, W303A *PIB2* ΔB::KAN, and W303A *PIB2* ΔhelE::KAN were generated using a similar approach, but by complete replacement of the His3MX6 cassette in the Δ*pib2*::HIS3 strain (PY_128). The replacement cassettes were generated by splicing by overlap extension using the appropriate *PIB2* construct amplified from pRS316 *S cer*. *PIB2* + UTRs (Varlakhanova et al., 2017), pRS316 *S cer*. *PIB2* ΔA + UTRs, pRS316 *S cer*. *PIB2* ΔB + UTRs, pRS316 *S cer*. *PIB2* ΔhelE + UTRs and a ADH1 terminator-kanMX6 fragment amplified from pFA6a-link-yEGFP-KanR (pKT0127) (Sheff and Thorn, 2004). After verification of correct reintroduction of the appropriate *PIB2* sequence by sequencing and loss of HIS prototrophy, the strains were further validated after backcrossing to W303α and resporulation.

Plasmids used in this work are listed in Supplemental Table S3. Pib2 mutant constructs were cloned by splicing by overlap extension (SOE) PCR at the site of the mutation using appropriate primers followed by Gibson assembly into the target vector. For Pib2 deletion constructs, the deleted regions were replaced with an AGAGA linker. N-terminal yEGFP-Pib2 constructs were generated with an N10 linker (NSSSNNNNNNNNNNLGIE). Targeting constructs were generated by amplification of the targeting protein from genomic DNA isolated from W303A/α diploids using the Yeast DNA Extraction kit (Thermo Scientific) or an existing plasmid. The targeting protein was then fused to the appropriate yEGFP-*PIB2* construct by SOE with a short linker between the targeting protein and yEGFP (GRRIPGLIN for *MVP1*-yEGFP constructs or GDGAGLIN for all others). These constructs also contain both the *PIB2* promoter (175 bp) and terminator (150 bp) and were fully assembled using Gibson assembly. All plasmids were verified by sequencing.

### Growth Analysis

Cells were grown overnight in YPD, SC, or SD with the appropriate dropout for plasmid maintenance. Cells were then diluted and regrown to mid-logarithmic phase (OD_600_ 0.5-0.8) in YPD at 30°C. Cells were diluted to 0.5 OD_600_/ml and 1:5 serial dilutions were made in water. Each dilution (2 μl) was spotted onto the indicated plates. Where relevant, cells were incubated with 200 ng/ml of rapamycin in YPD at 30°C for the indicated times. After several washes the cells were resuspended in fresh YPD and plated on YPD. All plates were incubated at 30°C and imaged on days 2 and 3.

### Preparation of yeast for microscopy

Cells were grown overnight in YPD or the appropriate SD medium. Cells were diluted in YPD and grown to mid-logarithmic phase. Vacuolar membranes were stained with 10 μM FM4-64 (Thermo Fisher Scientific) in YPD for 1 hour, followed by washing and incubation in SC media without dye for 1 hour. Cells were plated on No. 1.5 glass-bottomed cover dishes (MatTek Corporation, Ashland) treated with 15 μl 2 mg/ml concanavalin-A (Sigma-Aldrich).

### Microscopy and image analysis

Confocal images were acquired on a Nikon (Melville, NY) A1 confocal microscope, with a 100x Plan Apo 100x oil objective. NIS Elements imaging software was used to control image acquisition. Images were further processed using the Fiji distribution of ImageJ (Schindelin et al., 2012). GraphPad Prism was used for statistical analyses.

Vacuolar localization of yEGFP-Pib2 constructs was quantified using Fiji. For each cell, vacuolar membrane and cytosolic ROIs of equal area were determined and the fluorescence within those ROIs was measured. FM4-64 fluorescence was used to determine the location of the vacuolar membrane ROIs. Within each image, an average background fluorescence was determined and subtracted from the vacuolar and cytosolic intensity measurements. The localization was then expressed as a ratio of vacuolar membrane fluorescence to cytosolic fluorescence. For statistical analyses, a ROUT outliers test (Q = 0.1%) was used and data were further assessed by one-way ANOVA with Tukey multiple comparisons.

To determine expression levels of targeting constructs, cells were imaged in widefield using a Nikon Ti Microscope with an S Plan Fluor ELWD 20x objective. NIS Elements imaging software was used to control image acquisition. Images were processed in NIS Elements imaging software using a custom macro. GraphPad Prism was used for statistical analyses.

### Western Blotting

Protein extracts were prepared as previously described (Millen et al., 2009). Briefly, cells were lysed on ice by resuspension in 1 ml cold H_2_O supplemented with 150 μl 1.85 M NaOH and 7.5% (v/v) β-mercaptoethanol. After a 10-minute incubation on ice, the protein was precipitated by addition of 150 μl 50% (w/v) trichloroacetic acid. Pellets were washed twice with acetone, resuspended in 100 μl 1x SDS-PAGE buffer, and boiled for 5 minutes at 95°C. Primary antibodies were incubated overnight at 4°C and were as follows: anti-GFP (1:1000, ab290, Abcam), anti-PGK1 (1:1000, ab113687, Abcam), anti-Rps6 (1:1000, ab40820, Abcam), anti-phospho-Rps6 (1:1000, 4858, Cell Signaling Technology, Danvers). Secondary antibodies were incubated for 1 hour at room temperature and were as follows: IRDye 680RD goat ant-rabbit antibody (926-68171, Li-Cor, Lincoln) and IRDye 680D goat anti-mouse antibody (926-68070, Li-Cor). These were detected using the ChemiDoc MP Imaging System (Bio-Rad). Bands were integrated and quantified using Fiji.

### Size-exclusion chromatography coupled to multi-angle light scattering (SEC-MALS)

After filtration through a 0.22 μm cellulose acetate membrane, the Pib2 FYVE constructs and mutants were subjected to size-exclusion chromatography using a Superdex 75 10/300 equilibrated in SEC-MALS buffer (20 mM Tris/Cl, pH 7.4, 150 mM NaCl, 1.93 mM β-mercaptoethanol) at room temperature. 500 μl of each protein was loaded onto the column, at a concentration of 50 or 100 μM as indicated. The column was coupled to a static 18-angle light scattering detector (DAWN HELEOS-II) and a refractive index detector (Optilab T-rEX) (Wyatt Technology). Data were collected continuously at a flow rate of 0.3 ml/min, with the flow cells in the scattering and refractive index detectors set to 25°C. Data analysis was performed using the program Astra VII. Monomeric BSA (2.0 mg/ml) (Sigma) was used for data quality control.

### Isothermal Titration Calorimetry (ITC)

ITC was used to determine the thermodynamics of the interaction of the Pib2 FYVE domain (using Pib2 442-625) with inositol 1,3 bisphosphate (Ins(1,3)P2) or short-chain PI3P. The titrations were performed at 21°C using a PEAQ-ITC instrument (Malvern Analytical), in 20 mM Tris/Cl pH 7.4, 150 mM NaCl, 1.93 mM β-mercaptoethanol. 4 mM Ins(1,3)P2 or 2 mM diC4 PI3P was titrated into 100-150 μM protein. Integrated peaks for each titration were fit to a single site binding model using the MicroCal PEAQ-ITC software provided by the manufacturer.

## Notes

### Competing Interest Statement

The authors have declared no competing interest.

## References

Babst, M., Sato, T. K., Banta, L. M. and Emr, S. D. (1997). Endosomal transport function in yeast requires a novel AAA-type ATPase, Vps4p. EMBO J 16, 1820–31.

Babst, M., Wendland, B., Estepa, E. J. and Emr, S. D. (1998). The Vps4p AAA ATPase regulates membrane association of a Vps protein complex required for normal endosome function. EMBO J 17, 2982–93.

Belgareh-Touze, N., Avaro, S., Rouille, Y., Hoflack, B. and Haguenauer-Tsapis, R. (2002). Yeast Vps55p, a functional homolog of human obesity receptor gene-related protein, is involved in late endosome to vacuole trafficking. Mol Biol Cell 13, 1694–708.

Binda, M., Peli-Gulli, M. P., Bonfils, G., Panchaud, N., Urban, J., Sturgill, T. W., Loewith, R. and De Virgilio, C. (2009). The Vam6 GEF controls TORC1 by activating the EGO complex. Mol Cell 35, 563–73.

Bonfils, G., Jaquenoud, M., Bontron, S., Ostrowicz, C., Ungermann, C. and De Virgilio, C. (2012). Leucyl-tRNA synthetase controls TORC1 via the EGO complex. Mol Cell 46, 105–10.

Bridges, D., Fisher, K., Zolov, S. N., Xiong, T., Inoki, K., Weisman, L. S. and Saltiel, A. R. (2012). Rab5 proteins regulate activation and localization of target of rapamycin complex 1. J Biol Chem 287, 20913–21.

Brito, A. S., Soto Diaz, S., Van Vooren, P., Godard, P., Marini, A. M. and Boeckstaens, M. (2019). Pib2-Dependent Feedback Control of the TORC1 Signaling Network by the Npr1 Kinase. iScience 20, 415–433.

Burd, C. G. and Emr, S. D. (1998). Phosphatidylinositol(3)-phosphate signaling mediated by specific binding to RING FYVE domains. Mol Cell 2, 157–62.

Byfield, M. P., Murray, J. T. and Backer, J. M. (2005). hVps34 is a nutrient-regulated lipid kinase required for activation of p70 S6 kinase. J Biol Chem 280, 33076–82.

Chen, Z., Malia, P. C., Hatakeyama, R., Nicastro, R., Hu, Z., Peli-Gulli, M. P., Gao, J., Nishimura, T., Eskes, E., Stefan, C. J. et al. (2021). TORC1 Determines Fab1 Lipid Kinase Function at Signaling Endosomes and Vacuoles. Curr Biol 31, 297–309 e8.

Coonrod, E. M. and Stevens, T. H. (2010). The yeast vps class E mutants: the beginning of the molecular genetic analysis of multivesicular body biogenesis. Mol Biol Cell 21, 4057–60.

Dubouloz, F., Deloche, O., Wanke, V., Cameroni, E. and De Virgilio, C. (2005). The TOR and EGO protein complexes orchestrate microautophagy in yeast. Mol Cell 19, 15–26.

Edgar, R. C. (2004). MUSCLE: multiple sequence alignment with high accuracy and high throughput. Nucleic Acids Res 32, 1792–7.

Gonzalez, A. and Hall, M. N. (2017). Nutrient sensing and TOR signaling in yeast and mammals. EMBO J 36, 397–408.

Gonzalez, A., Shimobayashi, M., Eisenberg, T., Merle, D. A., Pendl, T., Hall, M. N. and Moustafa, T. (2015). TORC1 promotes phosphorylation of ribosomal protein S6 via the AGC kinase Ypk3 in Saccharomyces cerevisiae. PLoS One 10, e0120250.

Guan, J., Stromhaug, P. E., George, M. D., Habibzadegah-Tari, P., Bevan, A., Dunn, W. A., Jr. and Klionsky, D. J. (2001). Cvt18/Gsa12 is required for cytoplasm-to-vacuole transport, pexophagy, and autophagy in Saccharomyces cerevisiae and Pichia pastoris. Mol Biol Cell 12, 3821–38.

Hatakeyama, R. (2021). Pib2 as an Emerging Master Regulator of Yeast TORC1. Biomolecules 11.

Hatakeyama, R., Peli-Gulli, M. P., Hu, Z., Jaquenoud, M., Garcia Osuna, G. M., Sardu, A., Dengjel, J. and De Virgilio, C. (2019). Spatially Distinct Pools of TORC1 Balance Protein Homeostasis. Mol Cell 73, 325–338 e8.

Holt, L. J., Tuch, B. B., Villen, J., Johnson, A. D., Gygi, S. P. and Morgan, D. O. (2009). Global analysis of Cdk1 substrate phosphorylation sites provides insights into evolution. Science 325, 1682–6.

Ingrell, C. R., Miller, M. L., Jensen, O. N. and Blom, N. (2007). NetPhosYeast: prediction of protein phosphorylation sites in yeast. Bioinformatics 23, 895–7.

Jeong, J. H., Lee, K. H., Kim, Y. M., Kim, D. H., Oh, B. H. and Kim, Y. G. (2012). Crystal structure of the Gtr1p(GTP)-Gtr2p(GDP) protein complex reveals large structural rearrangements triggered by GTP-to-GDP conversion. J Biol Chem 287, 29648–53.

Jumper, J., Evans, R., Pritzel, A., Green, T., Figurnov, M., Ronneberger, O., Tunyasuvunakool, K., Bates, R., Zidek, A., Potapenko, A. et al. (2021). Highly accurate protein structure prediction with AlphaFold. Nature 596, 583–589.

Kamada, Y., Yoshino, K., Kondo, C., Kawamata, T., Oshiro, N., Yonezawa, K. and Ohsumi, Y. (2010). Tor directly controls the Atg1 kinase complex to regulate autophagy. Mol Cell Biol 30, 1049–58.

Kim, A. and Cunningham, K. W. (2015). A LAPF/phafin1-like protein regulates TORC1 and lysosomal membrane permeabilization in response to endoplasmic reticulum membrane stress. Mol Biol Cell 26, 4631–45.

Kira, S., Kumano, Y., Ukai, H., Takeda, E., Matsuura, A. and Noda, T. (2016). Dynamic relocation of the TORC1-Gtr1/2-Ego1/2/3 complex is regulated by Gtr1 and Gtr2. Mol Biol Cell 27, 382–96.

Kutateladze, T. G., Ogburn, K. D., Watson, W. T., de Beer, T., Emr, S. D., Burd, C. G. and Overduin, M. (1999). Phosphatidylinositol 3-phosphate recognition by the FYVE domain. Mol Cell 3, 805–11.

Loewith, R. and Hall, M. N. (2011). Target of rapamycin (TOR) in nutrient signaling and growth control. Genetics 189, 1177–201.

Loewith, R., Jacinto, E., Wullschleger, S., Lorberg, A., Crespo, J. L., Bonenfant, D., Oppliger, W., Jenoe, P. and Hall, M. N. (2002). Two TOR complexes, only one of which is rapamycin sensitive, have distinct roles in cell growth control. Mol Cell 10, 457–68.

MacGurn, J. A., Hsu, P. C., Smolka, M. B. and Emr, S. D. (2011). TORC1 regulates endocytosis via Npr1-mediated phosphoinhibition of a ubiquitin ligase adaptor. Cell 147, 1104–17.

Michel, A. H., Hatakeyama, R., Kimmig, P., Arter, M., Peter, M., Matos, J., De Virgilio, C. and Kornmann, B. (2017). Functional mapping of yeast genomes by saturated transposition. Elife 6.

Millen, J. I., Krick, R., Prick, T., Thumm, M. and Goldfarb, D. S. (2009). Measuring piecemeal microautophagy of the nucleus in Saccharomyces cerevisiae. Autophagy 5, 75–81.

Nakashima, N., Noguchi, E. and Nishimoto, T. (1999). Saccharomyces cerevisiae putative G protein, Gtr1p, which forms complexes with itself and a novel protein designated as Gtr2p, negatively regulates the Ran/Gsp1p G protein cycle through Gtr2p. Genetics 152, 853–67.

Nobukuni, T., Joaquin, M., Roccio, M., Dann, S. G., Kim, S. Y., Gulati, P., Byfield, M. P., Backer, J. M., Natt, F., Bos, J. L. et al. (2005). Amino acids mediate mTOR/raptor signaling through activation of class 3 phosphatidylinositol 3OH-kinase. Proc Natl Acad Sci U S A 102, 14238–43.

Panchaud, N., Peli-Gulli, M. P. and De Virgilio, C. (2013). Amino Acid Deprivation Inhibits TORC1 Through a GTPase-Activating Protein Complex for the Rag Family GTPase Gtr1. Science Signaling 6.

Parsons, A. B., Brost, R. L., Ding, H., Li, Z., Zhang, C., Sheikh, B., Brown, G. W., Kane, P. M., Hughes, T. R. and Boone, C. (2004). Integration of chemical-genetic and genetic interaction data links bioactive compounds to cellular target pathways. Nat Biotechnol 22, 62–9.

Peli-Gulli, M. P., Sardu, A., Panchaud, N., Raucci, S. and De Virgilio, C. (2015). Amino Acids Stimulate TORC1 through Lst4-Lst7, a GTPase-Activating Protein Complex for the Rag Family GTPase Gtr2. Cell Rep 13, 1–7.

Powis, K., Zhang, T., Panchaud, N., Wang, R., De Virgilio, C. and Ding, J. (2015). Crystal structure of the Ego1-Ego2-Ego3 complex and its role in promoting Rag GTPase-dependent TORC1 signaling. Cell Res 25, 1043–59.

Prouteau, M., Desfosses, A., Sieben, C., Bourgoint, C., Lydia Mozaffari, N., Demurtas, D., Mitra, A. K., Guichard, P., Manley, S. and Loewith, R. (2017). TORC1 organized in inhibited domains (TOROIDs) regulate TORC1 activity. Nature 550, 265–269.

Raymond, C. K., Howald-Stevenson, I., Vater, C. A. and Stevens, T. H. (1992). Morphological classification of the yeast vacuolar protein sorting mutants: evidence for a prevacuolar compartment in class E vps mutants. Mol Biol Cell 3, 1389–402.

Reinke, A., Anderson, S., McCaffery, J. M., Yates, J., 3rd, Aronova, S., Chu, S., Fairclough, S., Iverson, C., Wedaman, K. P. and Powers, T. (2004). TOR complex 1 includes a novel component, Tco89p (YPL180w), and cooperates with Ssd1p to maintain cellular integrity in Saccharomyces cerevisiae. J Biol Chem 279, 14752–62.

Robert, X. and Gouet, P. (2014). Deciphering key features in protein structures with the new ENDscript server. Nucleic Acids Res 42, W320–4.

Saxton, R. A. and Sabatini, D. M. (2017). mTOR Signaling in Growth, Metabolism, and Disease. Cell 168, 960–976.

Schindelin, J., Arganda-Carreras, I., Frise, E., Kaynig, V., Longair, M., Pietzsch, T., Preibisch, S., Rueden, C., Saalfeld, S., Schmid, B. et al. (2012). Fiji: an open-source platform for biological-image analysis. Nat Methods 9, 676–82.

Schrodinger, L. (2015). The PyMOL Molecular Graphics System, Version 2.4.0.

Sheff, M. A. and Thorn, K. S. (2004). Optimized cassettes for fluorescent protein tagging in Saccharomyces cerevisiae. Yeast 21, 661–70.

Shin, M. E., Ogburn, K. D., Varban, O. A., Gilbert, P. M. and Burd, C. G. (2001). FYVE domain targets Pib1p ubiquitin ligase to endosome and vacuolar membranes. J Biol Chem 276, 41388–93.

Shintani, T. and Klionsky, D. J. (2004). Cargo proteins facilitate the formation of transport vesicles in the cytoplasm to vacuole targeting pathway. J Biol Chem 279, 29889–94.

Stenmark, H., Aasland, R. and Driscoll, P. C. (2002). The phosphatidylinositol 3-phosphate-binding FYVE finger. FEBS Lett 513, 77–84.

Sturgill, T. W., Cohen, A., Diefenbacher, M., Trautwein, M., Martin, D. E. and Hall, M. N. (2008). TOR1 and TOR2 have distinct locations in live cells. Eukaryot Cell 7, 1819–30.

Subramanian, K., Dietrich, L. E., Hou, H., LaGrassa, T. J., Meiringer, C. T. and Ungermann, C. (2006). Palmitoylation determines the function of Vac8 at the yeast vacuole. J Cell Sci 119, 2477–85.

Sullivan, A., Wallace, R. L., Wellington, R., Luo, X. and Capaldi, A. P. (2019). Multilayered regulation of TORC1-body formation in budding yeast. Mol Biol Cell 30, 400–410.

Sun, D., Varlakhanova, N. V., Tornabene, B. A., Ramachandran, R., Zhang, P. and Ford, M. G. J. (2020). The cryo-EM structure of the SNX-BAR Mvp1 tetramer. Nat Commun 11, 1506.

Tanigawa, M. and Maeda, T. (2017). An in vitro TORC1 kinase assay that recapitulates the Gtr-independent glutamine-responsive TORC1 activation mechanism on yeast vacuoles. Mol Cell Biol.

Tanigawa, M., Yamamoto, K., Nagatoishi, S., Nagata, K., Noshiro, D., Noda, N. N., Tsumoto, K. and Maeda, T. (2021). A glutamine sensor that directly activates TORC1. Commun Biol 4, 1093.

Tarassov, K., Messier, V., Landry, C. R., Radinovic, S., Serna Molina, M. M., Shames, I., Malitskaya, Y., Vogel, J., Bussey, H. and Michnick, S. W. (2008). An in vivo map of the yeast protein interactome. Science 320, 1465–70.

Ukai, H., Araki, Y., Kira, S., Oikawa, Y., May, A. I. and Noda, T. (2018). Gtr/Ego-independent TORC1 activation is achieved through a glutamine-sensitive interaction with Pib2 on the vacuolar membrane. PLoS Genet 14, e1007334.

Urban, J., Soulard, A., Huber, A., Lippman, S., Mukhopadhyay, D., Deloche, O., Wanke, V., Anrather, D., Ammerer, G., Riezman, H. et al. (2007). Sch9 is a major target of TORC1 in Saccharomyces cerevisiae. Mol Cell 26, 663–74.

Varlakhanova, N. V., Mihalevic, M. J., Bernstein, K. A. and Ford, M. G. J. (2017). Pib2 and the EGO complex are both required for activation of TORC1. J Cell Sci 130, 3878–3890.

Wedaman, K. P., Reinke, A., Anderson, S., Yates, J., 3rd, McCaffery, J. M. and Powers, T. (2003). Tor kinases are in distinct membrane-associated protein complexes in Saccharomyces cerevisiae. Mol Biol Cell 14, 1204–20.

Yoon, M. S., Du, G., Backer, J. M., Frohman, M. A. and Chen, J. (2011). Class III PI-3-kinase activates phospholipase D in an amino acid-sensing mTORC1 pathway. J Cell Biol 195, 435–47.

